# ALFF response interaction with learning during feedback in individuals with multiple sclerosis

**DOI:** 10.1101/2022.04.27.489552

**Authors:** Ekaterina Dobryakova, Rakibul Hafiz, Olesya Iosipchuk, Joshua Sandry, Bharat Biswal

## Abstract

Amplitude of low-frequency fluctuations (ALFF) is defined as changes of BOLD signal during resting state (RS) brain activity. Previous studies identified differences in RS activation between healthy and multiple sclerosis (MS) participants. However, no research has investigated the relationship between ALFF and learning in MS. We thus examine this here. Twenty-five MS and 19 healthy participants performed a paired-associate word learning task where participants were presented with extrinsic or intrinsic performance feedback. Compared to healthy participants, MS participants showed higher local brain activation in the right thalamus. We also observed a positive correlation in the MS group between ALFF and extrinsic feedback within the left inferior frontal gyrus, and within the left superior temporal gyrus in association with intrinsic feedback. Healthy participants showed a positive correlation in the right fusiform gyrus between ALFF and extrinsic feedback. Findings suggest that while MS participants do not show a feedback learning impairment compared to the healthy participants, ALFF differences might suggest a general maladaptive pattern of task unrelated thalamic activation and adaptive activation in frontal and temporal regions. Results indicate that ALFF can be successfully used at capturing pathophysiological changes in local brain activation in MS in association with learning through feedback.

## Introduction

Multiple sclerosis (MS) is a central nervous system (CNS) disorder marked by demyelination, inflammation, and atrophy, resulting in long-term physical and cognitive impairments. Cognitive deficits affect between 43% and 70% of individuals with MS (Sumowski et al., 2018). Given that MS negatively affects the CNS diffusely, the number of investigations that use approaches that allow examination of the large-scale impact on CNS activity have been recently increasing. Analyzing brain activity at rest affords examination of brain activation while not being restricted by a behavioral task. However, it is still possible to examine brain activity at rest in association with a cognitive process through evaluating resting state (RS) fluctuations and behavioral performance in combination.

Numerous studies have documented altered RS activation in MS compared to healthy control (HC) participants. These investigations have shown that both adaptive and maladaptive brain reorganization in MS is associated with cognitive processes. For example, better cognitive performance (i.e., attention, information processing and working memory) in individuals with MS is associated with increased functional connectivity (FC) in the attention network (Loitfelder et al., 2012). Reduced resting-state functional connectivity (RSFC) of frontal regions in MS is associated with the severity of cognitive impairment (Bonavita et al., 2011; Parisi et al., 2014; Rocca et al., 2010a) and with the structural disruption of white matter tracts (Rocca et al., 2010b). At the same time, several studies (Hawellek et al., 2011; Schoonheim et al., 2015) report maladaptive patterns of RSFC along with associations between increased RSFC and worsened cognitive performance based on neuropsychological testing.

While the above studies examined RSFC, another method, amplitude of low frequency fluctuations (ALFF) (Zang et al., 2007), can also be applied to RS data. ALFF is a measure of the total power that lies within such low frequency bands (0.01 to 0.1 Hz specific to the current study). ALFF is a functional segregation method that provides local information based on RS-fMRI activity within specific brain regions. Biswal and colleagues originally demonstrated that the majority of the power in RS-fMRI signal is characterized by low frequency oscillations in the range 0 to 0.08 Hz (Biswal et al., 1995). The advantage of applying the ALFF method is that it is data driven and not influenced by any *a priori* hypothesis, contrary to seed-based functional connectivity and numerous other RS-fMRI analytical approaches. ALFF is also calculated on a voxel level, therefore, a whole brain map can be obtained, allowing for a broader inspection of the brain using information from local brain activation.

Given the positive characteristics of the ALFF methodology, there is still a paucity of studies utilizing this method in MS. Previous studies that compared ALFF activation between HC and MS participants reported lower ALFF values in the left hippocampus, limbic lobe, parahippocampal gyrus, temporal lobe, and caudate nucleus within the MS group. However, individuals with MS also show increased ALFF values in the limbic lobe, para-hippocampal gyrus, hippocampus, and amygdala (Gu et al., 2022). Another study that looked at activation patterns within the gray matter of the mirror neuron system, reported increased ALFF activation in the left inferior frontal gyrus (IFG). There was also a positive association between grey matter (GM) volume of inferior parietal lobule (IPL) and ALFF activity of the IFG. These differences were attributed to the differences in cognition among persons with MS due to variations in structural and functional neural abnormalities (Plata-Bello et al., 2018).

### Current study

In the current study, we explore differences in ALFF in RS data between MS and HC participants and whether ALFF response is associated with cognitive performance, specifically, learning feedback. Feedback learning is an important component of rehabilitation, including rehabilitation for persons with MS. Feedback can be either positive or negative and allows learners to modify their behavior to achieve a goal (Schlund & Pace, 1999) and is helpful in acquiring new skills. Positive feedback encourages an individual to continue the action being performed, while negative feedback encourages them to try a different action. Processing the information delivered through feedback is thus an important cognitive process that can either hamper or aid rehabilitation. Examining whether and how RS activation is related to feedback processing is an important step towards establishing the functional relevance of RS brain activation.

To examine the relationship between RS activation and feedback processing, we administered a paired-associate word learning task to HC and MS participants (Dobryakova et al., 2021; Dobryakova & Tricomi, 2013; Tricomi & Fiez, 2008). Participants were provided with two types of performance feedback: extrinsic feedback and intrinsic feedback. Extrinsic feedback represents an objective form of feedback, while intrinsic feedback represents a subjective form of feedback, where feedback value would vary from person to person. This direct feedback manipulation allowed us to not only investigate whether individuals with MS learn from feedback but also whether these two types of feedback influence learning differently.

## Materials and Methods

### Participants

Twenty-five persons with relapsing-remitting MS and twenty-two HC participants consented to participate in this study. Three out of 22 HC subjects included in the RS ALFF analysis did not have behavioral scores due to technical difficulties and therefore were removed from analyses that required behavioral data (see below). There were no group differences in age or education level (see Table 1 for demographic information). Participants were excluded if they reported a significant history of medical or psychiatric disorders, drug abuse, or learning disability. Participants with MS were at least 4 weeks post most recent exacerbation and use of steroids, benzodiazepines, or neuroleptics. The mean disease duration of those with MS was 14.12 and ranged from 4–27 years. The Institutional Review Board of Kessler Foundation approved the research protocol and all participants provided written consent to participate.

**Table 1.**
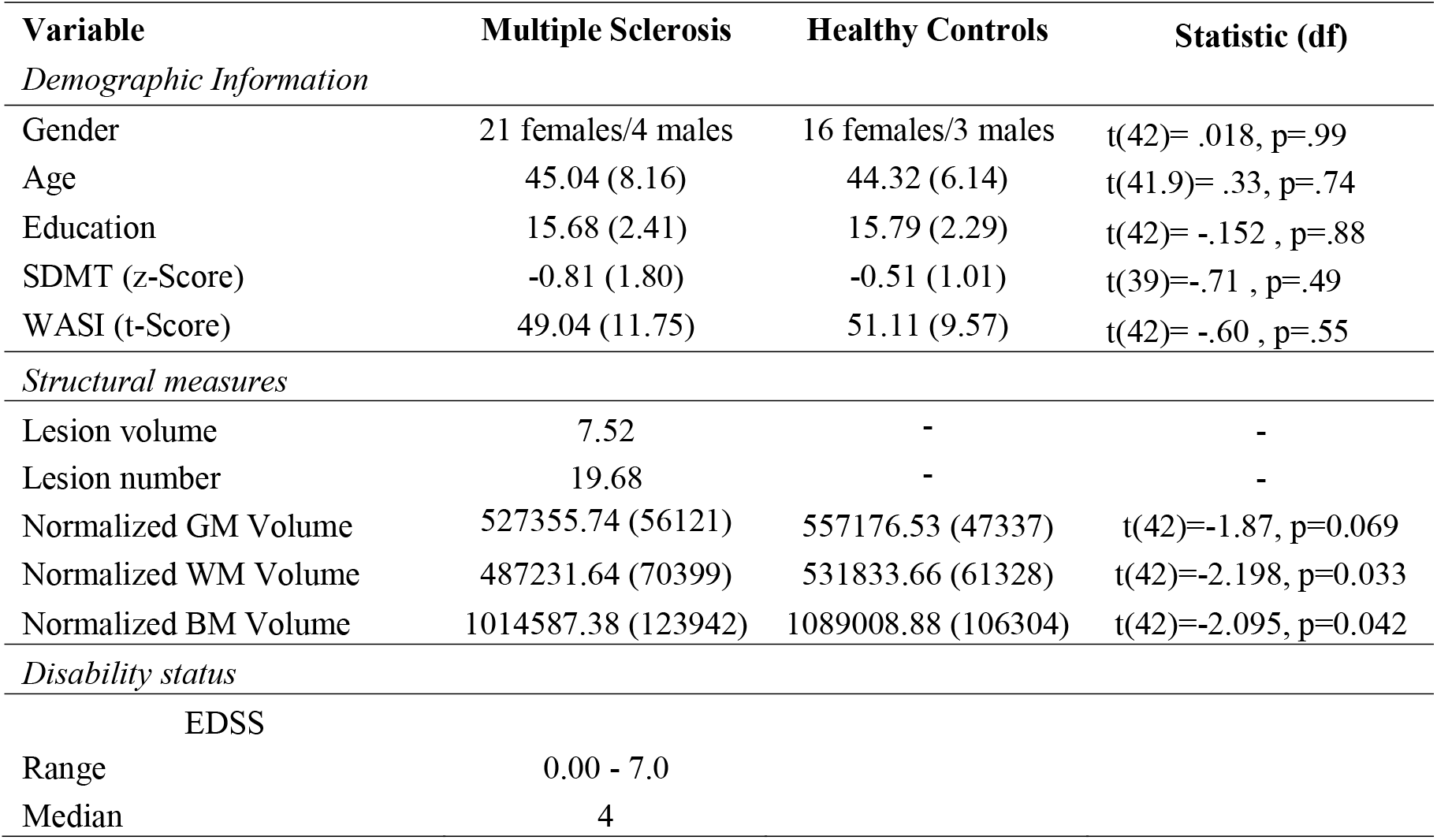
Data shows demographic information using means (standard deviation). Range and median are provided for EDSS. WM= white matter; GM= grey matter; BM = brain matter.

## Procedure

### Behavioral paradigm

Prior to beginning neuroimaging procedures, participants were told that they will undergo fMRI while engaging in a paired-associate word learning task. The learning task was adapted from previously published work and consist of the Study Phase, the Feedback Phase, and the Test Phase (Figure 1A) (Dobryakova et al., 2021; Dobryakova & Tricomi, 2013).

**Figure 1.**
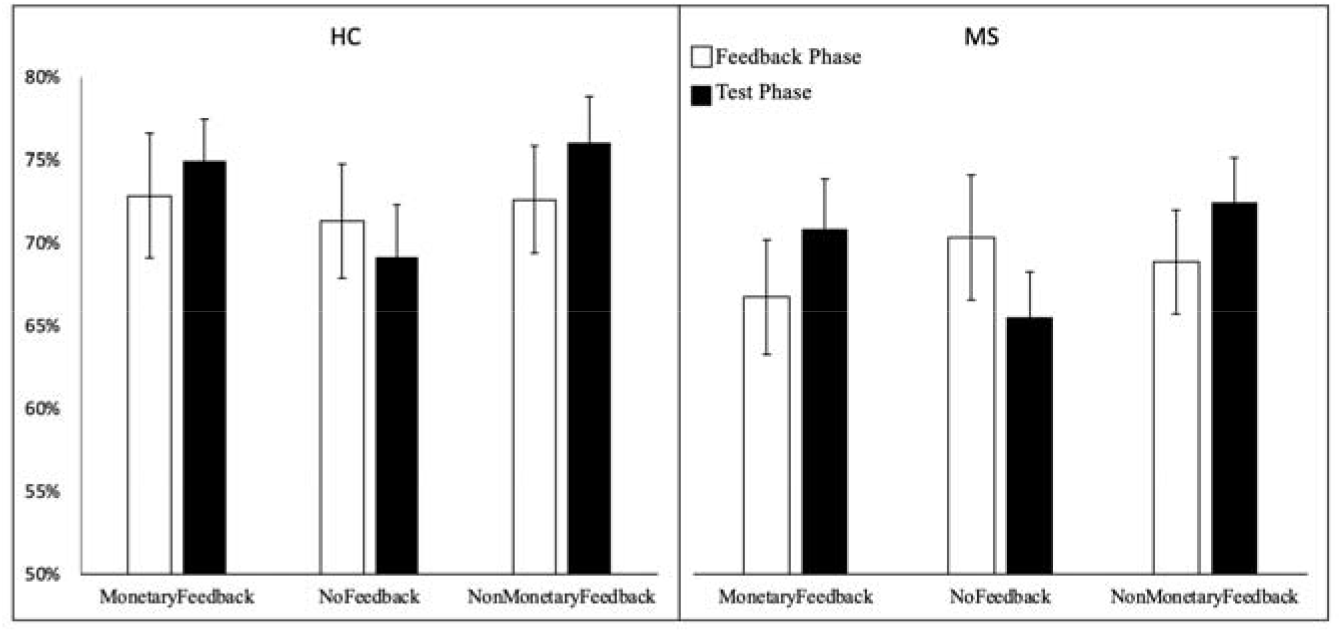
Accurancy for the three feedback conditions during the Feedback Phase and at Test Phase. Error bars represent standard error of the mean.

#### Study Phase

The Study Phase occurred outside of the scanner. During the Study Phase, participants learned word associations (180 trials). The format of experimental trials resembled a multiple-choice test. That is, on each trial of the Study Phase, participants were presented with three words: a target word and two-word options underneath. One of the options was highlighted in green, indicating to participants that this option is the correct match for the target word. Participants were instructed to memorize the target word and the associated highlighted option. The words used in the experiment contained 4–8 letters and 1–2 syllables, had Kucera-Francis frequencies of 20–650 words per million, and had high imageability ratings (score of over 400 according to the MRC database) (Coltheart, 1981). The words were matched for word length and frequency at the trial level. Words presented on the same trial were not semantically related, with a score of less than 0.2 on the Latent Semantic Analysis similarity matrix (Landauer et al., 1998). The words did not rhyme and did not begin with the same letter. Each trial was presented for 4 seconds to allow adequate time for memorization. Trials were presented in random order.

#### Feedback Phase

The words from the Study Phase were randomly assigned to one of three feedback conditions that were presented during the fMRI scan: Monetary Feedback (60 trials; MFB), Non-Monetary Feedback (60 trials; nMFB), and the No Feedback (60 trials; NFB). The conditions were presented randomly in blocks of 10 trials. Each trial stared with a jittered fixation point (1-5 s) that contained a label to inform participants of the condition they were presented with (Figure 1B). There were five blocks of each condition totaling 60 trials in each condition. The order of trials within each condition was randomized. During feedback trials, participants were presented with a cue screen that contained the target word with two words presented underneath. Participants selected the word that matched the target word based on what they remembered from the Study Phase. During the Monetary Feedback condition, if participants correctly matched the word to its paired associate target, they were presented with the feedback screen that had a green circle with a monetary gain amount written inside of it ($1.00); if they incorrectly matched the word to its target, they were presented with the feedback screen that had a red circle with a monetary loss amount written inside of it ($0.50). Participants were told that they will receive a bonus based on their performance. During the Non-Monetary Feedback condition, if participants correctly matched the cue word to its target, they were presented with the feedback screen that either had a green (if participants correctly matched the word to its target) or a red circle (if participants incorrectly matched the word to its target). During the No Feedback condition participants were presented with a blue circle. Each cue screen was presented for 4 seconds, and the feedback screen was presented for 1 second.

#### Test Phase

The final stage was performed outside the scanner. To test the impact of feedback on learning, participants were shown the same words as in previous phases in random order. Participants were presented with three words: a target word and two-word options underneath. The task was to choose one of the bottom words that was paired with the top word. Each time participants were presented with the choice screen for 4 seconds. Each trial was followed by the confidence question, which asked individuals to rate how confident they were with each response on a scale from 1 (complete guess) to 7 (completely sure).

At the end of their participation in the study, participants were debriefed and paid for participation ($150).

### Behavioral data analysis

Accuracy served as the main dependent measure. Accuracy data from the Feedback and Test Phases were analyzed with mixed effects ANOVA with Group (HC, MS) as a between subject factor and Phase (Feedback, Test) and Feedback condition (Monetary Feedback, Non-Monetary Feedback, No Feedback) as within subject factors. Additional analyses included a Phase X Feedback condition repeated measures ANOVA, computed independently for each group given the importance of inspecting clinical and non-clinical participants separately. These analyses were followed up by simple comparison analyses.

### fMRI data

#### Scan session

Participants received an MRI scan of the brain conducted on a Siemens Skyra 3T scanner. 32 slices of RS images were acquired using an EPI sequence (TR = 2,000 ms, TE = 30 ms, FOV = 220 mm, flip angle = 90°). 34 functional slices were collected using an EPI sequence (TR = 2,000 ms, TE = 30 ms, FOV = 204 mm, flip angle = 90°) tilted 30° from the AC-PC line to reduce signal drop-out in the VMPFC (Deichmann et al., 2003). A T1-weighted pulse sequence was used to collect structural images (TR = 2,100 ms, TE = 3.43 ms, flip angle = 9°, 1 × 1 × 1 mm^3^ voxels).

#### Pre-Processing

The pre-processing of RS data was performed using Statistical Parametric Mapping 12 (SPM12) toolbox (http://www.fil.ion.ucl.ac.uk/spm/) within the MATLAB environment (Mathworks Inc, Massachusetts, USA). After normalization, some modules from FSL *(FMRIB Analysis Group, Oxford, UK)* and AFNI (http://afni.nimh.nih.gov/afni) (Cox, 1996) were used for speed optimization, filtering and smoothing the data. The first five time points were removed to reduce transient scanner artifacts. Further data pre-preprocessing steps to reduce noise and increase signal-to-noise ratio are explained in detail below:

*Step1*: Motion correction for head movement using a least squared approach and 6 parameter (rigid body) spatial transformation with respect to the mean image of the scan was performed; 47 subjects (22 HC, 25 MS) were used for the following steps after eliminating 5 (2 HC, 3 MS) subjects with excessive head motion determined using framewise displacement (Power et al., 2012). A subject was eliminated if either the maximum framewise translation or rotation exceeded 1.5 mm, or the mean framewise displacement of translation exceeded 0.3 mm or, the mean displacement of rotation exceeded 0.2 mm. *Step2:* Co-registration of the mean functional image of each subject to their corresponding anatomical image was performed. *Step3:* Segmentation of anatomical images into gray matter, white matter (WM), and cerebrospinal fluid (CSF) tissue probability maps were conducted. *Step4:* Spatial normalization to the MNI space was done using deformation field vector obtained from the segmentation (in *Step 3*) of each subject and resampling to isotropic voxel size of 2 x 2 x 2 mm^3^. *Step5:* Regression of the first five principal components of WM and CSF time series extracted from voxels of WM and CSF probability maps thresholded > 0.98, along with other nuisance head motion noise using Friston’s 24-parameter model (6 head motion parameters + 6 previous time point motion parameters + 12 corresponding quadratic parameters)(Friston et al., 1996). *Step6:* Temporal filtering within the band 0.01 to 0.1 Hz and then finally, spatial smoothing was conducted on the data using a Gaussian kernel of 8 mm full width at half maximum (FWHM).

### Analysis

#### Local Brain Activation: ALFF

To investigate local brain activity and characterize spontaneous fluctuations of blood oxygen level dependent (BOLD) signal in the frequency domain, we computed voxelwise ALFF. The goals of this analysis are two folds: (i) identify where the differences in ALFF activity occur in MS brains compared to HCs and (ii) evaluate which brain regions demonstrate differences in interaction i.e., the slopes of the relationship between ALFF and the extrinsic and intrinsic feedback scores. Using AFNI’s ‘3dRSFC’, we generated whole brain ALFF maps and then converted them to Z-score maps for all subjects. These maps were subsequently concatenated group-wise in a General Linear Model (GLM) design for second level analysis in SPM12. The estimated model was used to run a parametric test using the unpaired two sample *t*-test design with age and gender added as covariates of no-interest. The resulting *t*-score statistical maps were thresholded at *t > 3.283* for an uncorrected height threshold of, *p_unc_ < 0.001.* The cluster-level extent threshold value, k_E_, was then identified from the SPM results table based on *family wise error (FWE)* correction for multiple comparisons at an alpha value threshold of *α = 0.05*. This was used to identify and visualize distinct clusters that demonstrated significant difference between groups at a corrected p-value threshold of *p_fwe_ < 0.05.*

For the second part of the analysis, a voxel-wise multiple linear regression approach was applied to evaluate the group level differences in interaction between ALFF and performance accuracy of the two behavioral feedback task scores (monetary and non-monetary feedback conditions). A GLM was generated in SPM12 by concatenating subject level Z-scored ALFF maps from HC and MS groups and a continuous covariate interaction model was set up for each feedback task score in turn. Age and gender of each subject were added as covariates of no interest in order to regress out and minimize the influence of confounding effects. The clusters demonstrating significant differences in interaction were identified and visualized, in the same manner as described above for the first analysis.

To assess and visualize the relationship between behavioral scores and ALFF, a graphical approach was adopted. Mean ALFF values from the clusters showing significant differences were extracted for each subject by first binarizing the thresholded statistical maps. These binary masks were used to extract the mean time series of the raw ALFF values for all subjects in HC and MS group. The values extracted from this step were plotted against the corresponding behavioral scores in a scatter plot and a line of best fit was generated for HC and MS group separately. The graphical analysis, general statistics and representation were performed using an inhouse script generated using RStudio (RStudio Team (2018). RStudio: Integrated Development for R. RStudio, Inc., Boston, MA URL http://www.rstudio.com/).

A significant difference in interaction in the direction HC > MS should be reflected by a positive slope of the ALFF and extrinsic/intrinsic feedback score relationship for HC and contrarily, a negative slope for the MS group. On the other hand, a significant difference in the MS > HC direction, should be reflected by a positive slope for MS and a negative slope for the HC group. A positive slope with a higher tangent is representative of a stronger positive correlation between mean local brain activation and the corresponding feedback score. On the contrary, a negative slope with higher tangent would represent a stronger negative correlation between these two variables.

#### Structural MRI data

To calculate total lesion volume and number of lesions in the MS sample, we used the Lesion Segmentation Toolbox for SPM. This toolbox uses FLAIR images to automatically segment T2 hyperintense lesions. SIENAX software was used to calculate normalized brain volume on T1-weighted lesions-filled images.

## Results

### Behavioral results

The 2×2×3 repeated measures ANOVA showed a significant phase by feedback condition interaction (F(2,42)=5.22, p=.007), as well as a significant main effect of feedback condition (F(2, 42)=21.77, p<.001). Additional analyses included a Phase X Feedback condition repeated measures ANOVA, computed independently for each group given the importance of inspecting clinical and non-clinical participants separately. In the MS group, we observed a significant phase by condition interaction (F(1, 24)= 7.89, p=.001). Analysis of the HC group also revealed a significant phase by condition interaction (F(1, 18)=3.57, p=.04) and a significant main effect of condition (F(1, 19)=7.05, p=.003).

Follow-up simple comparisons revealed that the MS group showed a significant improvement in accuracy from Feedback phase to Test Phase on the Monetary (t(24)=2.32, p=.03) and Non-Monetary (t(24)=2.1, p=.05) Feedback conditions and significant decrement in accuracy on the No Feedback condition (t(24)=2.1, p=.05). No significant differences were observed within the HC group (Figure 1).

### MRI results

#### Local Activation Differences: ALFF

We replicated previous findings of altered RS activation in MS using ALFF, revealing higher ALFF response in the right thalamus in the MS compared to the HC group (4, −14, 6; 14, −30, 2). Figure 2 shows the two distinct clusters and the extending subcortical regions demonstrating group level differences in ALFF. The clusters were generated after thresholding *t*-statistic maps at the *FWE* corrected cluster-level threshold according to SPM12’s two sample *t*-test results and visualized in AFNI. Peak t-score values were both observed at the right thalamus (Th) (see Table 2 for details).

**Figure 2.**
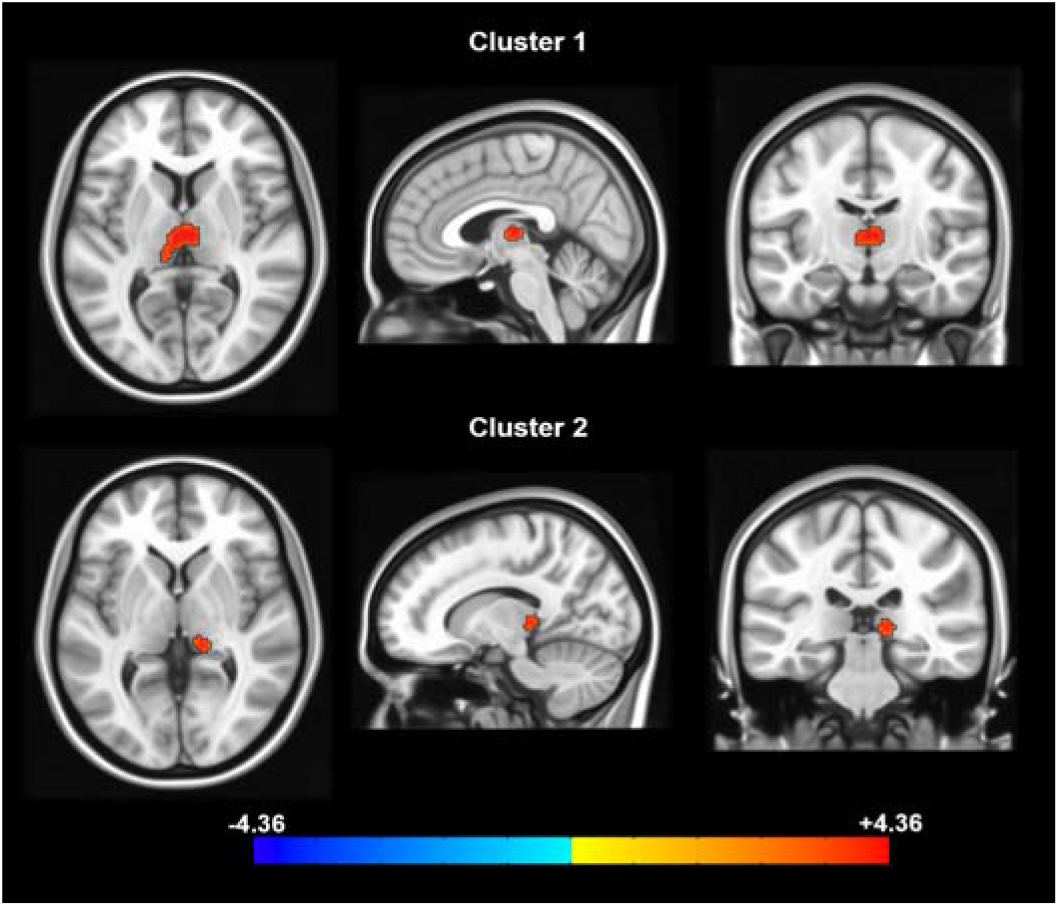
Significantly higher ALFF observed in the *right Thalamus* in MS group when compared to HC group. Two distinct clusters were obtained after cluster-based thresholding at height threshold *p_unc_ < 0.001* and *FWE* corrected at *p_FWE_ < 0.05* for multiple comparisons. The peak t-scores of clusters 1 (top row) and 2 (bottom row) were observed in the MNI coordinates: [x y z] = [4, −14 6] and [14 −30 2], with each cluster consisting of 229 and 53 voxels,

**Table 2.** Cluster details from the group level analyses of ALFF. The 1^st^ row provides details of group level ALFF differences observed between MS and HC groups in two distinct clusters. The 2^nd^ and 3^rd^ row provide cluster details pertaining to the interaction of ALFF with extrinsic and intrinsic in-scanner feedback scores, where MS group demonstrated higher effects compared to HC group. The final row also provides cluster details for the interaction analysis except, where HC group demonstrated higher effects compared to MS group for the extrinsic feedback, Keys: No. of. Cl. = Number of Clusters, MFB = Monetary Feedback, nMFB = non-Monetary Feedback.

| Analysis | Clusters | Region | Voxels | Peak X | Peak Y | Peak Z | Peak t |
| --- | --- | --- | --- | --- | --- | --- | --- |
| <u>ALFF</u> | 2 | Right Thalamus (Th) | 229 | 4 | -14 | 6 | 4.563 |
| MS > HC |  | Right Thalamus (Th) | 53 | 14 | -30 | 2 | 4.182 |
| <u>ALFF v. MFB</u> | 1 | Left Inferior Frontal | 335 | -48 | 18 | 6 | 4.480 |
| MS > HC |  | Gyrus (IFG) |  |  |  |  |  |
| <u>ALFF v. nMFB</u> | 1 | Left Temporal Pole | 271 | -60 | 10 | -2 | 4.411 |
| MS > HC |  | (TP) |  |  |  |  |  |
| <u>ALFF v. MFB</u> | 1 | Right Fusiform Gyrus | 101 | 30 | -28 | -20 | 4.197 |
| HC > MS |  | (FuG) |  |  |  |  |  |

#### ALFF and Behavioral Scores

When performance accuracy from the task was regressed on RS data, the MS group exhibited greater ALFF response compared to the HCs in *frontal* and *temporal* regions. Three out of 22 HC subjects included in the local activation differences analysis above did not have behavioral scores due to technical difficulties and therefore were removed from the continuous covariate interaction analysis, thus, this section shows results from 19 HC and 25 MS. ALFF response interaction with learning from extrinsic feedback (monetary feedback condition [MFB]) was higher in the MS group in the *left inferior frontal gyrus (IFG)* (−48, 18, 6), while interaction with learning from intrinsic feedback (non-monetary feedback condition [nMFB]) was higher in the MS group in the *left superior temporal gyrus (STG)* (−60, 10, −2).

Figure 3A shows brain regions in the MS group that demonstrated a significantly higher interaction between ALFF and extrinsic feedback scores compared to the HC group. This is reflected in the two graphs at the bottom of Figure 3A. The graph on the left represents a negative correlation between mean ALFF values and extrinsic (MFB) feedback scores in the HC group. On the other hand, the graph on the bottom right shows positive correlation between mean ALFF values and extrinsic (MFB) feedback scores in MS group. Similarly, Figure 3B shows brain regions and corresponding graphs for the MS group, demonstrating significantly higher interaction between ALFF and intrinsic (nMFB) feedback scores *in the left temporal pole (TP) and superior temporal gyrus (STG) (see* Table 2 for details).

**Figure 3.**
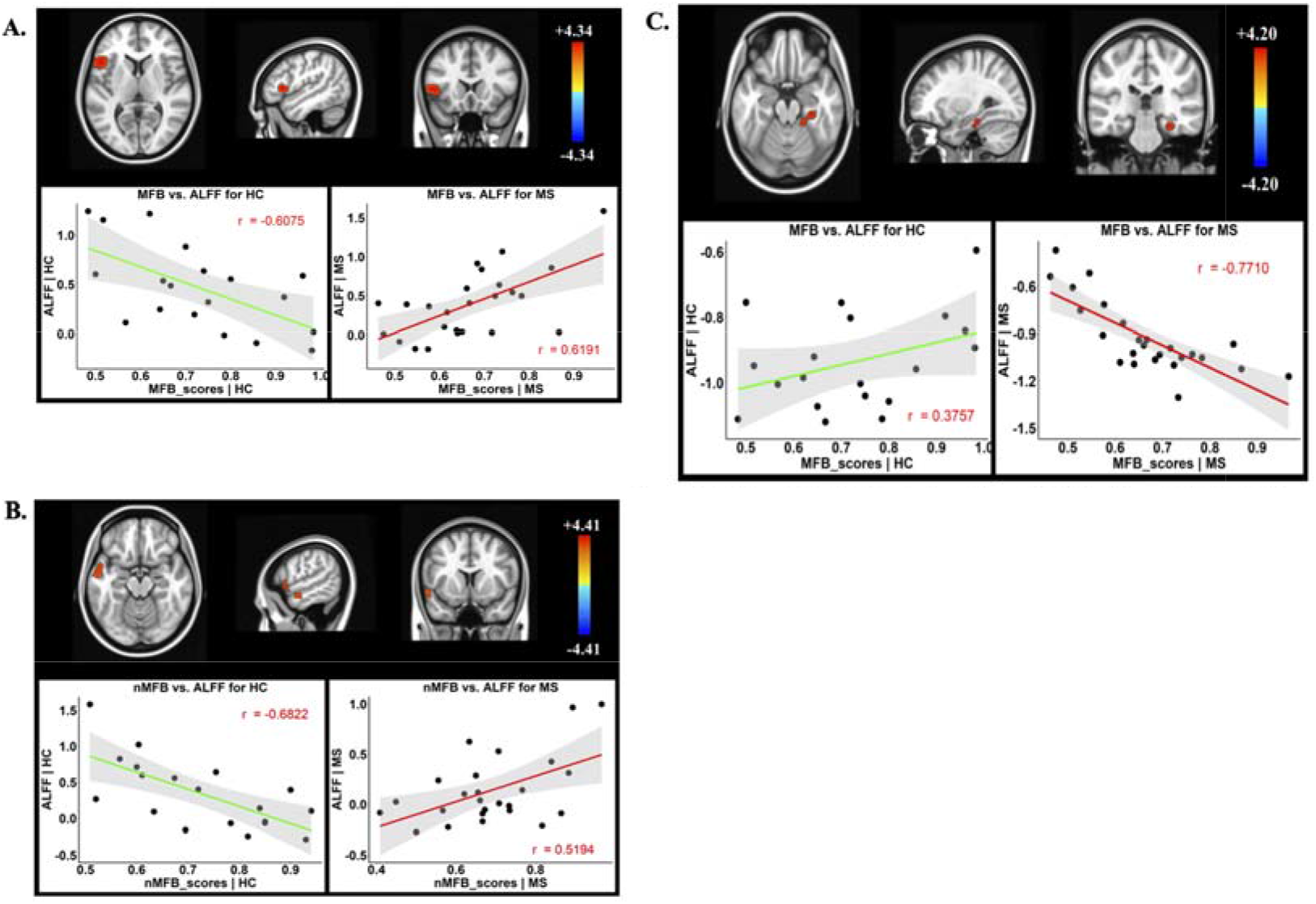
MS > HC | Higher interaction of ALFF with extrinsic (MFB) and intrinsic (nMFB) feedback scores in MS compared to HC group. A. Top row: Brain regions demonstrating significant differences in interaction of ALFF and extrinsic feedback score in the *IFC.* Bottom left: Negative correlation between mean ALFF values extracted from the 1FG cluster and extrinsic feedback scores from the HC group. Bottom Right: Positive correlation between mean ALFF values and extrinsic feedback scores in MS group. B. Similar to descriptions in A., except, for intrinsic feedback scores, which demonstrated significantly higher interaction with ALFF in MS compared to HC group in the *left TP/STG* cluster. C. HC > MS ļ Higher interaction of ALFF with extrinsic (MFB) feedback scores in HC compared to MS group. Top: Brain regions demonstrating higher interaction of ALFF and extrinsic feedback score in HCs in the *right FuG.* Bottom Left: Positive correlation between mean ALFF values and extrinsic feedback scores in the HC group. Bottom Right: Negative correlation between mean ALFF values and extrinsic feedback scores in the MS group. Keys: MFB = Monetary Feedback [Extrinsic Feedback], nMFB = non-Monetary Feedback [Intrinsic Feedback]. The y-axis represents the mean ALFF values of the cluster for each subject. The x-axis represents the corresponding feedback scores from same subjects. The green/red line represents the line of best fit and the grey area represents the 95% confidence interval of the line of best fit.

While the brain regions that demonstrate higher interaction are left-lateralized in MS group, only the *right fusiform gyrus* (*FuG*) (30, −28, −20) showed greater ALFF response interaction in HCs with learning from extrinsic (MFB) feedback. Figure 3C depicts significantly higher interaction of local brain activation and extrinsic feedback (MFB) scores in HC compared to MS. The graphs at the bottom of Figure 3C provide a visual interpretation of this relationship. The bottom left figure in Figure 3C shows positive correlation of mean ALFF values from the significant cluster with extrinsic monetary feedback scores in HC group. The bottom right graph of Figure 3C shows a negative correlation of mean ALFF scores with extrinsic feedback scores, indicating lower local activation response interaction in MS group. The peak t-score was observed at the *FuG* with more cluster details provided in Table 2.

These findings suggest that while persons with MS do not show a behavioral impairment of learning through feedback compared to the HC group, differences in ALFF response might suggest a general maladaptive pattern of task unrelated *thalamic* ALFF activation and adaptive activation during rest in the MS group in *frontal* and *temporal* regions.

## Discussion

The goal of the current study was to investigate local resting state activation differences between relapsing-remitting MS and HC participants using ALFF analysis and to further assess the differences in interaction between ALFF and learning from extrinsic and intrinsic feedback between these groups. While no behavioral differences in learning from feedback were observed between groups, analysis of ALFF demonstrated both adaptive and maladaptive differences in the MS group compared to the HC group. We discuss these findings in the context of the current literature below.

Results from the ALFF analysis indicate that the MS group demonstrates higher local brain activity in the right thalamus that further extends bilaterally to the left thalamus (see Figure 2). Several studies have reported thalamo-cortical abnormalities in MS subjects, but these investigations primarily focused on understanding FC alterations. For example, Lin et al. (2019) used whole brain seed-based FC of four thalamic sub-regions from the anterior, lateral and medial locations and showed that FC changes tend to be nuclei selective. Later, a more comprehensive study reported FC abnormalities in several other regions – middle frontal gyrus, sensorimotor network, precuneus, insula, and cerebellum (Hidalgo de la Cruz et al., 2018). Both stronger and weaker FC was observed corresponding to different sub-nuclei of the thalamus. Other studies based on parcellation of thalamic regions (d’Ambrosio et al., 2017) and seed-based approach have shown various thalamic abnormalities across MS cohorts (van Geest et al., 2017). However, FC analysis reflect temporal dependencies between two or more spatial regions, while ALFF reflects local neural activity, which can be more informative of the pathophysiological changes in the MS brain. Previous studies have shown increased ALFF in the bilateral thalami of MS patients (Y. Liu, Liang, Duan, Jia, Wang, et al., 2011; Zhou et al., 2014). Zhou et al. (2014) further showed a strong correlation between ALFF and fractional anisotropy (FA) in the left thalamus, thereby, indicating a relationship between functional and structural changes in these subjects. Our local activation analysis using ALFF align with these findings, further highlighting the role of thalamic integrity.

More importantly, using a continuous covariate interaction model, we aimed to further assess how ALFF alterations correlate with learning from extrinsic and intrinsic feedback. This second part of the analysis highlights the novelty of this study. Our results show that compared to HCs during both extrinsic and intrinsic feedback learning, MS subjects demonstrate a stronger correlation with ALFF, primarily in the left IFG and left TP. On the other hand, for the extrinsic feedback, HCs demonstrate a stronger correlation in the FuG within the right hemisphere. A recent study reported decreased ALFF in the IFG among relapsing remitting MS subjects (Plata-Bello et al., 2018). However, that study included patients that had milder disability (median expanded disability status scale [EDSS]=2.0) than the current sample (median EDSS=4.0). Moreover, the ALFF analysis from Plata-Bello et al. (2018) was limited to the ROIs within the mirror neuron system and not applied to the whole brain, as is the case in the current study. It is also important to note that their results showed positive correlation between ALFF of left IFG and gray matter volume (GMV) from the left inferior parietal lobule (IPL) in MS subjects. Interestingly, when we assessed the interaction between ALFF and extrinsic feedback, we also found positive correlation between ALFF and extrinsic feedback performance in the left IFG in the MS group, whereas HCs demonstrated a negative correlation. The clinical significance of this outcome still needs further investigation to identify if there is a structural relationship such as white matter tract damage or morphological changes in GMV due to disease progression.

For the intrinsic feedback, MS group demonstrated positive correlation with ALFF compared to negative correlation in the HC group within the left TP. One study had shown increased ALFF among MS subjects in the right temporal lobe, particularly, in the STG, and further demonstrated that these alterations have a significant correlation with disability (through EDSS scores) (Y. Liu, Liang, Duan, Jia, Yu, et al., 2011). On the contrary, Y. Liu et al. (2015) reported decreased ALFF in MS subjects both in the left and right middle temporal gyrus (MTG). However, the MS subjects in their study also had simple spinal cord involvement, which may contribute towards such bilateral decrease in the MTG (Y. Liu et al., 2015). Our results show a positive correlation between ALFF and intrinsic feedback scores in the left temporal lobe, particularly, in the left TP in the MS group, compared to negative correlation in HC group. While our results indicate increased ALFF in the temporal lobe, similar to Y. Liu et al. (2011) and in contrast to Y. Liu et al. (2015), they also suggest that alterations in different spatial locations vary depending on a cognitive task through which interaction with brain derived quantities is being estimated; comorbid injuries may also ‘co-alter’ local neural activity. Adding on to the alterations observed in the temporal lobe in association with extrinsic feedback, the HC group demonstrated a positive correlation between the temporal lobe and ALFF compared to a negative correlation between the right FuG and ALFF in the MS group. H. Liu et al. (2016) reported increased ALFF in the right FuG among relapsing remitting MS patients when compared to HCs. This inconsistency could indicate that while local activation may increase in MS patients, ALFF’s interaction during learning through extrinsic feedback could be anticorrelated in selective spatial locations, such as the right FuG, in this case. This may also point towards some maladaptive pathophysiological changes that may have caused local neural activity to alter significantly in MS patients when compared to HCs. Therefore, in future research, a more comprehensive study design can be developed where both structural and functional changes are observed longitudinally, along with clinical evaluation and cognitive task performance, to delineate the pathophysiological alterations that may elicit such brain-behavior relationships.

## Conclusion

In the current investigation, we showed that individuals with MS exhibited altered ALFF activation in the thalamus compared to healthy individuals. Further, we also observed altered ALFF activation as it relates to learning form extrinsic and intrinsic feedback. While we did not observe between group difference in learning from extrinsic and intrinsic feedback, our findings show that differences in ALFF response might suggest a general maladaptive pattern of task unrelated thalamic ALFF activation and adaptive activation during rest in the MS group. Future studies should more comprehensively examine both structural and functional changes in association with learning though feedback to delineate the pathophysiological alterations that may elicit observed brain-behavior relationships.

## Acknowledgements

The authors would like to thank Kessler Foundation for support and participants who contributed their time to participate in this study.

